# Single nucleotide switches confer bacteriophage resistance to *Pseudomonas protegen*s

**DOI:** 10.1101/2024.11.12.622978

**Authors:** Jordan Vacheron, Clara M. Heiman, Christoph Keel

**Affiliations:** Department of Fundamental Microbiology, University of Lausanne, Lausanne, Switzerland

## Abstract

Phage therapy offers a promising strategy against bacterial pathogens in medicine and agriculture, but the rise of phage-resistant bacteria presents a significant challenge to its sustainability. Here, we used an environmental model bacterium, *Pseudomonas protegens* CHA0, to investigate phage resistance mechanisms in laboratory conditions through genomic analysis of four phage-resistant variants (C2, C4, C17, C18). Whole-genome sequencing revealed frequent deletions, insertions, and single nucleotide substitutions, particularly in genes encoding enzymes involved in cell surface modifications. The T428P mutation in AlgC, a phosphoglucomutase, and the P229T substitution in YkcC, a glycosyltransferase, each conferred resistance by potentially altering phage receptor accessibility while preserving bacterial fitness. These findings suggest that subtle mutations in surface-modifying enzymes enable phage resistance by modifying phage-receptor accessibility while maintaining bacterial growth, highlighting adaptive mechanisms in bacterial phage interactions. Further structural analysis is needed to fully understand these mutation-driven resistance mechanisms.

## Introduction

The alarming rise of antimicrobial resistance (AMR) is one of the most pressing challenges of modern medicine and environmental microbiology (Larsson and Flach 2022; Darby *et al*. 2023). Various strategies have been proposed to combat this problem, some of which include destabilizing bacterial membranes to enhance antibiotic uptake (Lin *et al*. 2015) or inhibiting efflux pumps to reduce antibiotic release and consequently increase antibiotic concentration in bacterial cells (Alav, Bavro and Blair 2022). While these strategies aim to make current antibiotics more efficient by thwarting resistance mechanisms, a biologically driven approach is resurfacing. This approach permits to target specific bacteria without collateral damage to beneficial commensal microorganisms by relying on the specific interaction between bacteria and their natural predators, bacteriophages. Bacteriophages, or phages and their derivatives (e.g. tailocins or lysins) currently represent a promising alternative to combat AMR, as evidenced by recent cases, such as the remission of a patient previously infected with multidrug-resistant *Pseudomonas aeruginosa* (Köhler *et al*. 2023). However, relying on phages solely, even in combination with antibiotics, often falls short of ensuring pathogen eradication. As against antibiotics, bacteria have evolved multiple mechanisms to resist phage attacks, leading to a parallel arms race between bacteria and antibiotics as well as phages.

One prominent strategy employed by bacteria to resist phage infection involves the utilization of the CRISPR-Cas system for adaptive immunity, storing and recalling phage DNA fragments (Bernheim and Sorek 2020). In addition to this strategy, bacteria can also produce anti-phage proteins that disrupt different stages of the life cycle of their viral predator. Bacteria can also modify their cell surface decorations that serve as phage receptors. For instance, bacteria can alter or mask these receptors, making it challenging for phages to recognize and bind to their target cells (Bernheim and Sorek 2020). The interplay between bacteriophages and bacteria, shaped by these diverse resistance mechanisms, highlights the complexity of the microbial warfare that unfolds in diverse environments, including clinical and agricultural settings.

Phage therapy has also shown promising results in the biological control of bacterial plant pathogens (Wang *et al*. 2019). However, bacterial inoculants used to improve the growth and health of agricultural crops are also challenged by naturally occurring phages. Indeed, a lytic bacteriophage was discovered in the rhizosphere of cucumber plants that specifically targets the plant-beneficial and entomopathogenic strain *Pseudomonas protegens* CHA0 (Keel *et al*. 2002). This phage belonging to the *Zobelviridae* family and designated ΦGP100 (Vacheron *et al*. 2023), had a significant effect in reducing the abundance of CHA0 in the rhizosphere of cucumber plants. This reduction resulted in the loss of the plant protection against a root pathogenic oomycete (Keel *et al*. 2002). The molecular interaction between the phage ΦGP100 and CHA0 was investigated, and the long O-polysaccharide (O-PS) of the lipopolysaccharide (LPS) of CHA0 was found to be responsible for bacterial susceptibility (Vacheron *et al*. 2023). During the 2002 experiment, several spontaneous phage-resistant variants were isolated (Keel *et al*. 2002). These variants appeared within the clear halo two days following phage exposure of CHA0 on plates. As a proof of concept, these CHA0 variants were not affected by the phage for their root colonization, and they provided the plant pathogen suppressive effect.

Here, we have sequenced four of these phage-resistant CHA0 variants to pinpoint the specific mutations that confer resistance in a laboratory setting. Our results revealed that these mutations occured in genes responsible for cell surface decorations and resulted from single nucleotide changes leading to amino acid substitutions. One mutation substitutes a threonine for a proline in AlgC, a critical enzyme involved in lipopolysaccharide biosynthesis, potentially impacting the structure of the enzyme without abolishing its function. Another mutation affects YkcC, a glycosyltransferase encoded within the OBC3 (O-PS biosynthesis cluster) gene cluster involved in the production of the long O-antigen of CHA0, potentially altering cell surface structures targeted by the phage ΦGP100. These specific amino acid substitutions confer resistance while maintaining bacterial fitness *in vitro*. Our findings highlight how single nucleotide substitutions can modulate cell surface components, thereby conferring an adaptive advantage against phage infection.

## Materials and methods

### Genome sequencing and variant calling analysis

Four phage ΦGP100-resistant bacterial variants of *Pseudomonas protegens* CHA0 (C2, C4, C17 and C18) were obtained within the clear lytic halo, after two to three days of exposure of a bacterial lawn of CHA0 to the phage (Keel *et al*. 2002). The four CHA0 variants were cultured overnight in nutrient yeast broth (NYB) and their genomic DNA was extracted using the MagAttract high-molecular-weight (HMW) DNA kit (Qiagen). DNA samples were then sheared in Covaris g-TUBEs to obtain fragments with an average length of 10 ⍰kb. The PacBio SMRTbell template preparation kit v1.0 was used to prepare the library. The resulting library was size selected on a BluePippin system (Sage Science, Inc.) for molecules larger than 7⍰kb. The library was multiplexed and sequenced using one single-molecule real-time (SMRT) cell and a Sequel system (movie length, 600 min). Genome assembly was performed using the RS_HGAP_Assembly.4 protocol in SMRT Link v6.0. Variant calling analysis was performed using Snippy (version 4.3.6. Seeman) to identify genomic differences that could explain the emergence of the phage resistance in the genomes of the four ΦGP100-resistant variants of CHA0 (C2, C4, C17 and C18). The complete genome of *P. protegens* CHA0 (accession number LS999205, Smits et al. 2019) was used as reference for these analyses. Graph representations were generated using the R package circlize (Gu *et al*. 2014) in RStudio (R version 4.4.0).

### Deletion mutants and site-directed mutagenesis

Target gene deletions were performed following the procedures previously outlined in Vacheron et *al*. 2019. Site-directed mutagenesis was realized as previously described in Kupferschmied et al. 2014. Briefly, the entire genes of interest (either *ykcC* or *algC*) were cloned into the suicide plasmid pEMG, resulting in plasmids designated as pME11180 and pME11181, respectively (**Table S1 and S2, Supporting Information**). PCR amplification was then performed, using these plasmids as templates and sets of primers designed to introduce specific mutations to subsequently replace the desired amino acid residues. Following PCR amplification, the plasmid templates used were digested with DnpI for 1 h at 37°C. PCR products were purified using the QIAquick PCR Purification Kit (Qiagen, Deutschland). The purified PCR products were then electroporated into *E. coli* DH5α λpir cells. Selection of *E. coli* that successfully recircularized the vectors was performed on lysogeny broth (LB) agar supplemented with kanamycin (25 µg mL^-1^). Once these vectors were obtained (**Table S1, Supporting Information**), they were electroporated into CHA0 for double recombination as described previously by Kupferschmied et al. 2014. All constructed plasmids and mutant strains were controlled by DNA sequencing (Microsynth AG).

### Validation of the phage resistance phenotypes

The sensitivity of the CHA0 wild type and the different variants and mutants to the phage ΦGP100 was tested using the double layer agar method. Briefly, 50 µL of overnight bacterial cultures were mixed with 4⍰mL of LB soft agar (7.5 % of agar) and poured onto a nutrient agar plate. Serial dilutions of the phage suspension were spotted (4 µL) onto the bacterial lawn. All the plates were incubated overnight at 25⍰°C.

### Analysis of amino-acid conservation and protein structure visualization

To determine if the amino acid substitutions observed in the natural phage-resistant variants of CHA0 affected the conserved regions of the proteins through bacterial evolution, we aligned the amino acid sequences of AlgC and YkcC orthologs detected in *Pseudomonas* and more distantly related bacterial genomes. Amino-acid sequences were obtained by Blastp and aligned using ClustalO (Sievers and Higgins 2014). Amino-acid conservation was characterized for each residue position by calculating the Shenkin divergence score (Shenkin, Erman and Mastrandrea 1991) and visualized using JalView (Troshin *et al*. 2018). The structures of AlgC and YkcC were predicted using Alphafold 3 (Abramson *et al*. 2024) and visualized with ChimeraX v1.8 (Meng *et al*. 2023).

### Protein purification

The wild-type and variant proteins of AlgC were produced and purified using standard techniques. Briefly, the wild-type and mutant coding sequences for *algC* were cloned into a pET-based expression vector under control of a T7 promoter (**Table S1, Supporting Information**). A His-tag was added at the N-terminus to enable pulldown after overexpression in *Escherichia coli*. The resulting expression vectors were transformed into chemically competent BL21(DE3) cells and clones were selected on LB plates containing 50 µg mL^-1^ of kanamycin. Colonies were then inoculated into 10 mL of TB (terrific broth) medium and grown to an OD_600nm_ of 0.5 at 37 °C. The cultures were then cooled down to 18 °C and expression was induced by the addition of 0.4 mM of IPTG and incubated overnight (16 h). To collect samples for ‘total protein’ contents of the bacteria, 60 µL of each culture was spun down and the pellet was resuspended in 50 µL of 1x SDS loading dye (50 mM Tris-HCl pH 6.8, 5 % beta-mercaptoethanol, 2 % SDS, 0.1 % bromophenol blue, 10 % glycerol). The suspension was then heated to 95 °C for 10 min and centrifuged at 14,000 rpm for 10 min to pellet cell debris. Eight µL of the resulting supernatants were loaded onto SDS-PAGE gels. The remainder of the overnight culture (10 mL) was centrifuged at 5000 rpm for 10 min. The supernatants were discarded, and the pellets were resuspended in 2 mL of lysis buffer (50 mM Tris pH 7.5, 300 mM NaCl, 10 % glycerol, 25 mM imidazole). The cell suspensions were then sonicated to lyse the cells, and the lysate was clarified by centrifugation at 15,000 rpm at 4 °C. The supernatant was incubated with magnetic His-affinity resin (HisMag Sepharose, Cytiva, USA) for 1 h at 4 °C on a rotating wheel. Then the beads were collected on a magnetic rack and washed twice with 1 mL of lysis buffer. Finally, the bound material was eluted by incubation with 30 µL of elution buffer (lysis buffer supplemented with 500 mM imidazole). The supernatant was mixed with an equal volume of 2 x SDS loading dye (100 mM Tris-HCl pH 6.8, 10 % beta-mercaptoethanol, 4 % SDS, 0.2 % bromophenol blue, 20 % glycerol), heated to 95 °C for 5 min, and 8 µL of this solution was loaded onto an SDS-PAGE gel. After the run, the gel was fixed using a fixing solution (50 % ethanol, 10 % acetic acid) and then stained with Coomassie Brilliant Blue (50 % methanol, 10 % acetic acid, 2 mg mL^1^ Coomassie Brilliant Blue R-350).

### Bacterial growth monitoring

To test for differences in the growth of the variants and mutants with the wild type CHA0, bacterial suspensions obtained from an overnight NYB culture were restarted (1:100 v/v) in fresh medium until they reached an OD_600nm_ of 0.5. Then 5 µL of a bacterial suspension from an OD_600nm_ of 1 were inoculated in each well of the 96-well plate containing NYB rich medium (i.e., final OD_600nm_ of 0.05). The bacterial growth of the different strains was monitored every 10 min for 24 h using a Synergy H1 plate reader (BioTek, USA). Three independent replicates were performed. Growth parameters such as the carrying capacity, the growth rate, the generation time and the area under the curve were calculated using the R package GrowthCurver (Sprouffske and Wagner 2016). Statistical differences were determined by ANOVA coupled with Tukey’s HSD test, after assessing the dataset for normality and homoscedasticity.

### Lipopolysaccharide extraction and visualization on SDS-PAGE

LPS were examined using previously established methods (Davis and Goldberg 2012; Heiman *et al*. 2022). After treatment with proteinase K, the samples were subjected to sodium dodecyl sulfate polyacrylamide gel electrophoresis (SDS-PAGE) on either an 8 %-12 % gradient or 12 % acrylamide gel and visualized by silver staining. The PageRuler Plus Prestained Protein Ladders (Thermo Scientific) were used as molecular weight standards. The LPS extraction and visualization were performed four times independently.

## Results and discussion

### Mutation inventory within phage-resistant variants of Pseudomonas protegens CHA0

The four bacterial variants resistant to the phage ΦGP100 were originally isolated during double agar experiments, where small colonies emerged within the lysis zone of the phage (Keel *et al*. 2002). Their resistance to the phage was confirmed in the present study by conducting double-layer agar experiments, using each of the isolated variants for the second layer (**Fig. 1**).

**Fig 1:**
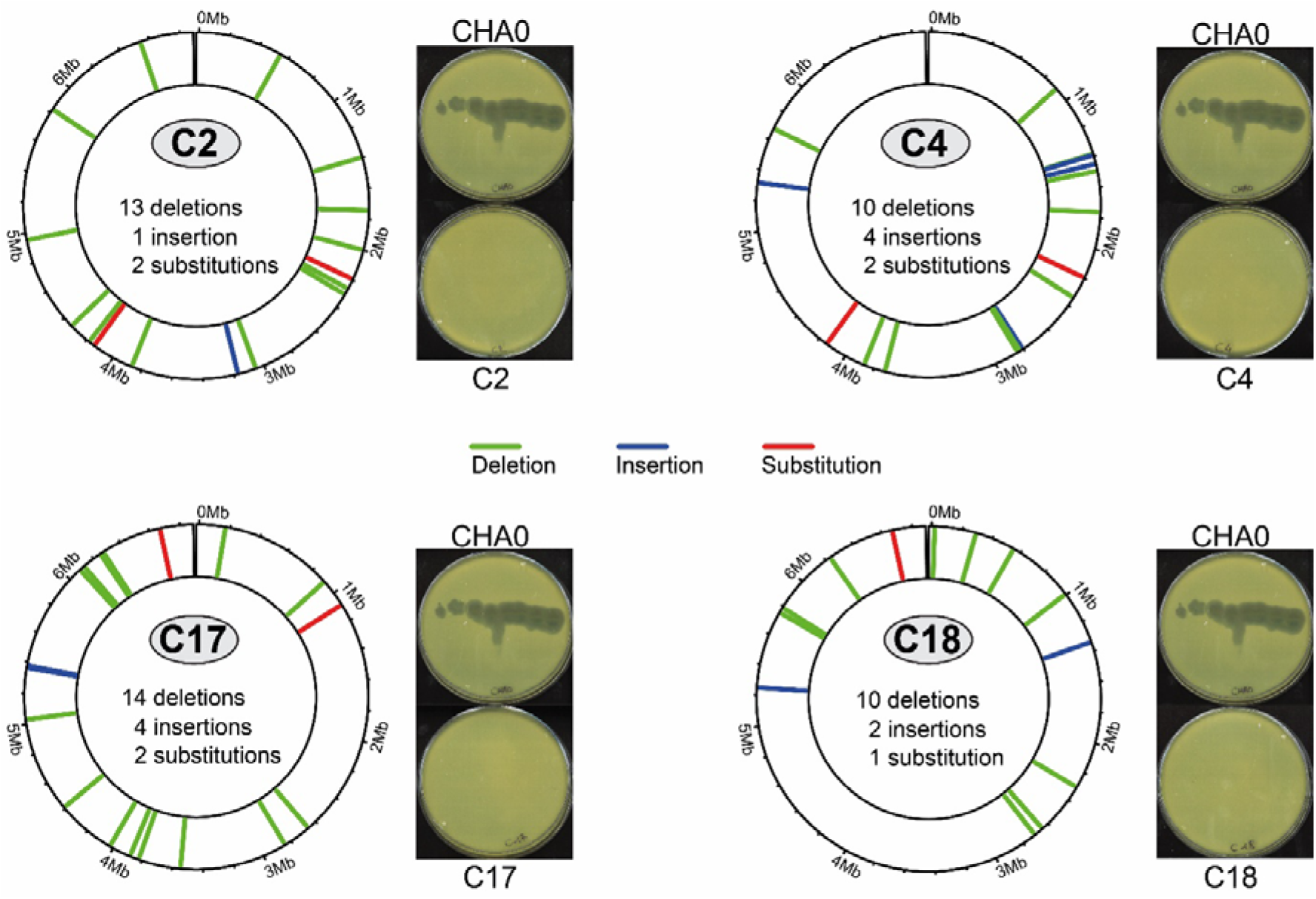
Variant calling analysis reveals the distribution of mutations in the genomes of the phage-resistant variants C2, C4, C17 and C18 of *Pseudomonas protegens* CHA0. Overview of the distribution of mutations occurring within the coding sequences for the different phage-resistant variants. The genomic locations of the different mutations are represented by red, blue, and green lines for substitution, insertion, and deletion mutations, respectively. On the side of each circular genome representation, sensitivity assays on plate show the phage resistance of the different variants compared to the wild type CHA0.

To identify the genetic basis of phage resistance in the four different CHA0 variants (C2, C4, C17, and C18), long-read sequencing (PacBio) was used to identify mutations. The genome sequences of the phage-resistant variants were compared with the complete genome sequence of the CHA0 wild type (LS999205.1). Genomic analysis of the phage-resistant bacterial variants revealed a complex landscape of mutations, with both shared patterns and unique differences among the strains (**Fig. 1; Tables S3 to S6, Supporting Information**). Deletions in coding sequences emerged as the predominant type of mutation, occurring in all variants and ranging from 10 to 14 events per genome. In addition to deletions, insertions and substitutions were also identified, but less frequently. Furthermore, between 14 and 21 mutations were detected in intergenic regions for each variant, with several of these mutations being common among variants. The wild-type genome, sequenced in 2019 (Smits *et al*. 2019), has been continuously used and shared across laboratories since 2002 (date of isolation of the different phage-resistant variants). By comparison, the high number of mutations observed in the variants, both within intergenic and coding regions, may be attributed to their long-term storage, as they were kept frozen for over 20 years (Keel et al. 2002) without further use. Laboratory-adapted strains are well documented for *Pseudomonas aeruginosa* PAO1, which has accumulated mutations over successive passages and across different laboratory environments, resulting in several cases in the modification of several bacterial strain phenotypes (Klockgether *et al*. 2010; Chandler *et al*. 2019). This laboratory-derived strain effect may explain a certain number of the sequence differences observed in the present study. In addition, the CHA0 reference genome, which is considered a complete genome, was sequenced using short-read sequencing technology, whereas the variants in this study were sequenced using long-read technology, which is known to be more accurate because it can span longer repetitive genomic regions (Espinosa *et al*. 2024).

Despite the sequence differences attributable to laboratory manipulation of the reference strain CHA0 and the use of a different sequencing technology, several of the identified mutations in the variants match those previously discovered by high-density transposon insertion sequencing (Tn-Seq) analysis (**Fig. 1; Tables S3 to S6, Supporting Information;** Vacheron et al. 2023). This makes them the main target for trying to understand which mutations cause the emergence of phage resistance in these CHA0 variants. Among them, a substitution in AlgC observed in the genomes of variants C17 and C18, and a substitution in a glycosyltransferase (YkcC) within the gene cluster for the long O-polysaccharide in the genomes of variants C2 and C4, are potential candidates (**Tables S3 to S6, Supporting Information)**. Since these cell surface decorations were previously identified as key players in phage sensitivity using a Tn-Seq approach (Vacheron *et al*. 2023), we aimed to confirm that these single nucleotide changes observed in the phage-resistant variants specifically confer resistance to the phage ΦGP100 and to understand their impact on the affected proteins.

## Single nucleotide substitution in the vicinity of the active site of AlgC confers phage resistance

AlgC is a bisubstrate mutase that transfers phosphate from position 6 to position 1 on glucose and mannose (Coyne *et al*. 1994; Olvera *et al*. 1999; Feng *et al*. 2023). This enzyme plays a crucial role in carbohydrate metabolism by catalyzing the interconversion of glucose-1P and glucose-6-P or mannose-1P and mannose-6P, which are important intermediates in the biosynthesis of glycoconjugates, such as LPS in bacteria (Coyne *et al*. 1994). Spontaneous mutants C17 and C18, resistant to phage ΦGP100 (**Fig. 1**), were found to harbor a single nucleotide substitution (C1279G) in *algC*, resulting in a T428P amino acid change (**Fig. 2A**).

**Fig 2:**
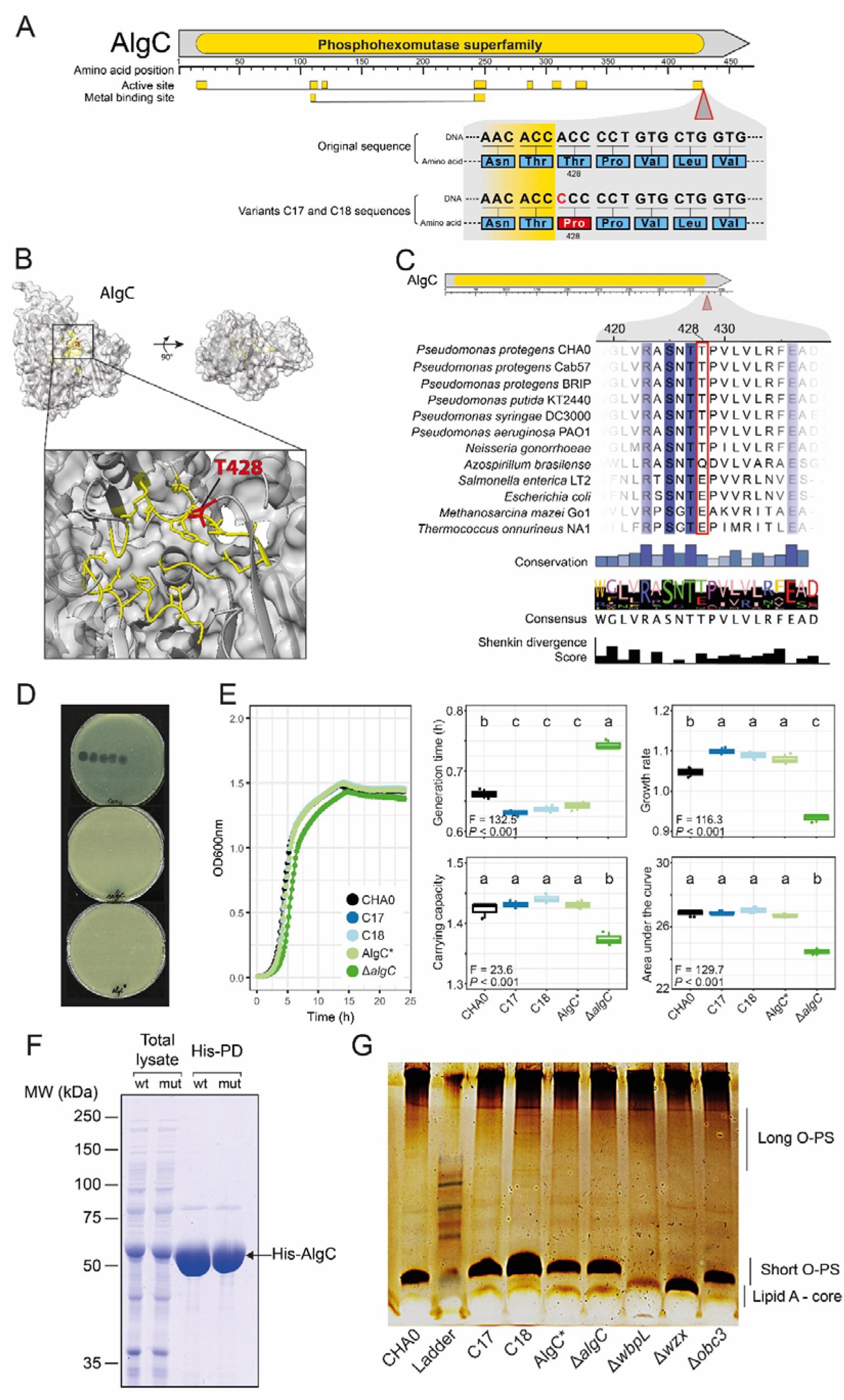
Characterization of the spontaneous AlgC mutation and its impact on phage sensitivity and bacterial fitness. (**A**) The amino acid change present in variants C17 and C18 substitutes threonine 428 for a proline. Active sites and metal-binding sites are highlighted in yellow. (**B**) Localization of the proline-substituted threonine in the AlgC protein. The amino acids constituting the different active sites in panel A are highlighted in yellow. The amino acid highlighted in red corresponds to the one substituted with a proline in the natural phage-resistant variants C17 and C18. (**C**) Amino acid sequence comparisons for different strains showing the conservation of the different amino acids in the site of interest. Amino acid conservation, consensus sequence and Shenkin divergence score are shown below. (**D**) Bacterial lawns of CHA0, the site-directed mutant AlgC* and the deletion mutant Δ*algC* exposed to serial dilutions of the phage GP100. (**E**) Impact of the AlgC mutation on the bacterial growth of CHA0, its variants and derivatives. Statistical differences were assessed by ANOVA coupled with the HSD Tukey test. Significant differences are indicated with letters a, b and of His-tagged AlgC proteins. Lane 1: Total lysate from wild-type (wt) cells. Lane 2: Total lysate from AlgC T428P mutant cells. Lane 3: His-tag pull-down (His-PD) from wt cells. Lane 4: His-PD from mutant cells. The position of the His-tagged AlgC protein is indicated by the arrow. (**G**) Lipopolysaccharide (LPS) profiles of CHA0 wt, the phage-resistant variants C17 and C18, AlgC* (site-directed mutant P428), Δ*algC*, as well as previously characterized control strains (Kupferschmied *et al*. 2016) Δ*wbpL* (truncated core LPS, no short or long O-polysaccharides (O-PS)), Δ*wzx* (no short O-PS), and Δ*obc3* (no long O-PS). SDS-PAGE on LPS extracts was performed using a 12% acrylamide gel and components were visualized by silver staining. The extraction and visualization of LPS was performed four times, other gel photos are available in **Fig. S1, Supporting Information**).

The T428 residue is located near the active site of AlgC (**Fig. 2A and 2B**). Although distant in the linear amino acid sequence, the predicted protein folding (Alphafold prediction) brings these predicted active sites into proximity, forming a functional pocket (**Fig. 2B**). This specific folding likely contributes to the catalytic activity of AlgC by positioning key residues, such as T428, in close interaction with the active site. Sequence alignment reveals that the T428 residue in AlgC is conserved across species within the *Pseudomonas* genus and *Neisseria gonorrhoeae* while it can be different in other bacterial species (**Fig. 2C**). Although there are variations in the residues observed at this position in other species, none of these species carry a proline at this position, as seen in the C17 and C18 mutants. On the one hand, this variation in orthologs across different bacterial lineages suggests a degree of functional flexibility in the active site of AlgC, which could have implications for its role in glycoconjugate biosynthesis, phage resistance, and other cellular processes. On the other hand, replacing threonine with a proline could have significant implications on the enzyme functionality, as prolines are known for their distinctive properties that can significantly affect protein structure (Bajaj *et al*. 2007; Melnikov *et al*. 2016). Indeed, unlike most amino acids, proline possesses a cyclic structure, forming a rigid ring, which restricts its flexibility. This characteristic makes proline stand out in the context of protein structures, as it introduces kinks or bends in the polypeptide chain (Bajaj *et al*. 2007; Melnikov *et al*. 2016).

The complete deletion of *algC* in the genome of CHA0 confers resistance to phage ΦGP100 (**Fig. 1**), confirming our previously identified involvement of AlgC in the sensitivity of CHA0 to the phage using a Tn-seq approach (Vacheron *et al*. 2023). To infer whether the amino acid substitution detected in the phage-resistant variants C17 and C18 is responsible for the emergence of the phage resistance phenotype, we used site-directed mutagenesis to generate this substitution in the wild-type CHA0 (strain AlgC*). Using double agar assays, we confirmed that the replacement of T428 with a proline is sufficient to confer the phage resistance phenotype to CHA0 (**Fig. 2D**).

Given the important role of AlgC in glucose metabolism, we assessed the impact of the mutation in this gene on the bacterial fitness by looking at bacterial growth parameters (**Fig. 2E**). Our results demonstrate that the C17 and C18 variants, as well as the site-directed mutant, exhibit shorter generation times and higher growth rates compared to the wild type, suggesting that the mutation confers a fitness advantage to these strains. In addition, the deletion mutant, which lacks the entire *algC* gene, showed a significant fitness impairment, emphasizing the essential role of AlgC in supporting optimal bacterial growth under the conditions tested. These findings indicate that the modified AlgC in the phage-resistant CHA0 variants retains its full or partial functionality.

To determine if the mutation affects AlgC folding and/or protein stability, we His-tagged both the wild-type and variant proteins and purified them following heterologous expression in *E. coli* (**Fig. 2F**). The bands corresponding to wild-type and variant AlgC appeared identical in size and intensity (**Fig. 2F**), indicating that the variant protein is produced at comparable levels. These findings suggest that the mutation does not alter the physico-chemical properties of AlgC, such as solubility, confirming that the variant remains stable and soluble.

A previous study using an *algC* mutant showed the importance of this enzyme for the core LPS formation (Coyne *et al*. 1994). To investigate potential modifications in LPS composition in phage-resistant variants and mutants, we analyzed their LPS profiles by electrophoresis (**Fig. 2G, Fig. S1, Supporting Information**). We analyzed the LPS profiles of different mutants to help us to clarify and identify the LPS modifications occurring in the naturally phage-resistant variants. The *wbpL* mutant, used as a control strain impaired in core LPS formation, had no detectable short or long O-PS bands as previously reported (Kupferschmied *et al*. 2016). Analysis of the *wzx* mutant confirmed the absence of short O-PS, consistent with its previously reported phenotype, while it remained sensitive to phage ΦGP100 (Kupferschmied *et al*. 2016). The *algC* deletion mutant showed a core lipid A and short O-polysaccharide (O-PS) pattern similar to the wild type. This finding contrasts with previous studies reporting that *algC* deletions cause core LPS truncation (Coyne *et al*. 1994). The preserved LPS core in the *algC* mutant might suggest functional redundancy in fully assembled cell surface decorations, despite growth impairment of the strain (**Fig. 2E**). However, no *algC* ortholog was detected in the genome of CHA0. In contrast, the phage-resistant variants C17 and C18 displayed a more pronounced short O-PS band in their LPS profiles. However, the long O-PS was barely visible, making it difficult to draw any definitive conclusions about its presence or modifications (**Fig. 2G, Fig. S1, Supporting Information**). We suggest that an increased capping with short O-PS along with a potential modification of the long O-PS (not visible on the SDS-PAGE gels) could interfere with phage adsorption, conferring the resistance to the phage ΦGP100. The amino acid substitution in AlgC of the variants C17 and C18 did not impair their fitness: growth rates were slightly enhanced, and generation times reduced (**Fig. 2E**).

In summary, the mutation in *algC* identified in the phage-resistant variants C17 and C18 enables the production of a stable and functional enzyme, despite the substitution of threonine with proline. Moreover, this mutation does not impair bacterial fitness. Additionally, the core O-polysaccharide remains detectable in the variants, indicating that AlgC retains its functionality. Altogether, we suggest that this mutation may have altered the role of AlgC in LPS production, resulting in modified cell surface decorations that hinders phage infection in these bacterial strains.

### An amino acid switch in YkcC glycosyltransferase confers phage resistance to CHA0

The *ykcC* gene encodes a glycosyltransferase within the O-antigen biosynthetic cluster OBC3, which is involved in the biosynthesis of long O-polysaccharides (O-PS) in CHA0 (Kupferschmied *et al*. 2016). The long O-PS was previously identified as the main receptor for infection of CHA0 by the phage ΦGP100 (Vacheron *et al*. 2023).

In the phage-resistant variants C2 and C4, a mutation substitutes a proline (P) with a threonine (T) at position 229 (**Fig. 3A**). To test the impact of this substitution, we introduced the T229P mutation into the wild-type strain via site-directed mutagenesis, generating the YkcC* variant. Phage susceptibility assays confirmed that this single mutation as well as the total deletion of *ykcC* confer resistance to phage ΦGP100, as shown by the absence of phage infection in the double agar assay (**Fig. 3B**).

**Fig 3:**
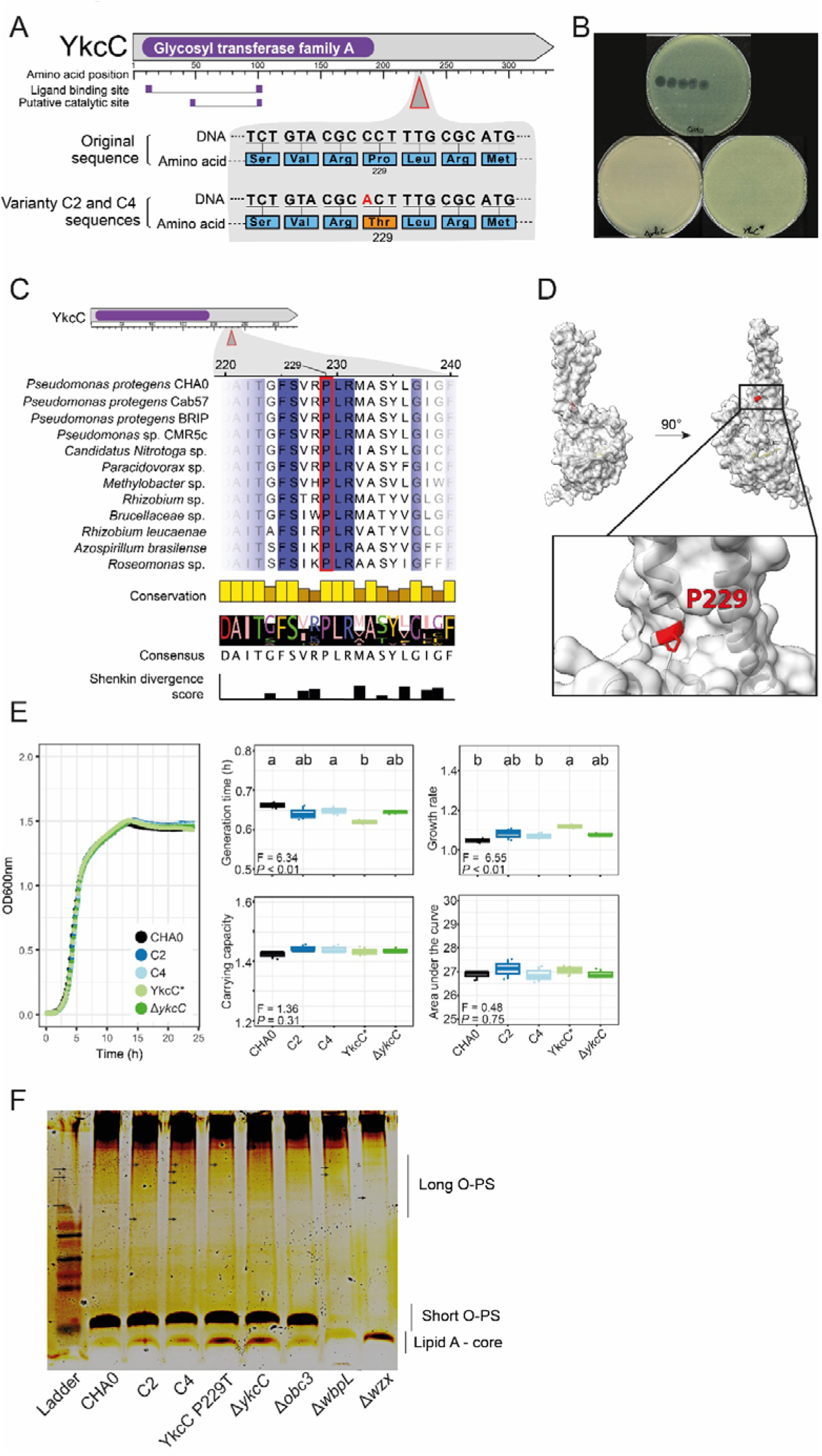
Characterization of the spontaneous YkcC mutation and its impact on phage sensitivity and bacterial fitness. (**A**) The amino acid change present in variants C2 and C4 substitutes proline 229 for a threonine. Active sites and metal-binding sites are highlighted in purple. (**B**) Bacterial lawns of CHA0, the site-directed mutant YkcC* and the deletion mutant Δ*ykcC* exposed to serial dilutions of the phage GP100. (**C**) Amino acid sequence comparisons for different strains showing the conservation of the different amino acids in the site of interest. Amino acid conservation, consensus sequence and Shenkin divergence score are shown below. (**D**) Localization of the threonine-substituted proline in the YkcC protein. The amino acids constituting the different active sites in panel A are highlighted in yellow. The amino acid highlighted in red corresponds to the one substituted with a proline in the natural phage-resistant variants C2 and C4. (**E**) Impact of the YkcC mutation on the bacterial growth of CHA0, its variants and derivatives. Statistical differences were assessed by ANOVA coupled with the HSD Tukey test. Significant differences are indicated with letters a, b and c. Three independent biological experiments were performed, each with three technical replicates. (**F**) Lipopolysaccharide (LPS) profile of CHA0 wild-type, the phage-resistant variants C2 and C4, YkcC* (site-directed mutant T229), Δ*algC*, as well as previously characterized control strains (Kupferschmied *et al*. 2016) Δ*wbpL* (truncated core LPS, no short or long O-polysaccharides (O-PS)), Δ*wzx* (no short O-PS), Δ*obc3*(no long O-PS). SDS-PAGE on LPS extracts was performed using a 12% acrylamide gel and components were visualized by silver staining. The extraction and visualization of LPS was performed four times, other gel photos are available in **Fig. S2, Supporting Information**).

The proline at position 229 is highly conserved among orthologs of this glycosyltransferase, suggesting a potential key role in maintaining the structure and function of this enzyme (**Fig. 3C**). Although this substituted amino acid is distant from the predicted catalytic site of YkcC (**Fig. 3D**), the switch from proline to threonine could induce a substantial structural alteration potentially affecting the overall folding of the protein and its function. To better characterize the impact of this substitution on the production, folding of the protein, a His-tag was added at the N-terminus of the wildtype and the variant copy of YkcC to enable protein purification after overexpression in *Escherichia coli*. Unfortunately, the purification of YkcC was unsuccessful, potentially due to the low solubility of this protein.

We then wanted to obtain phenotypic information that could explain the role of this mutation in YkcC. First, growth assays showed that the mutation did not affect the growth rate of phage-resistant variants C2, C4, or the site-directed mutant YkcC* (**Fig. 3E**). Next, we analyzed the LPS structure of these strains. Surprisingly, the LPS profile of C2, C4, YkcC*, and the *ykcC* deletion mutant resembled that of the wild type, with no noticeable changes in the long O-antigen region, which YkcC is expected to influence (**Fig. 3F, Fig. S2, Supporting Information**). Since YkcC is part of the OBC3 gene cluster involved in long O-PS production (Kupferschmied *et al*. 2016), we anticipated a modification here; however, no distinct differences were observed.

One hypothesis for this unexpected result is that the P229T mutation might alter the structure of YkcC in a subtle way that affects its interaction with phage ΦGP100 without visibly impacting O-PS biosynthesis. This structural shift could mask phage receptors by influencing O-antigen composition at a molecular level, thus preventing efficient phage binding and conferring resistance. We are also aware that the silver staining method employed here to visualize LPS may not provide sufficient resolution to properly observe changes in the long O-PS of CHA0. Further structural studies are needed to fully understand the impact of this mutation on YkcC functionality and phage resistance.

## Conclusion

In this study, we identified specific genetic mutations in *Pseudomonas protegens* CHA0 that confer resistance to phage ΦGP100. Through whole-genome sequencing of phage-resistant variants, we discovered single nucleotide substitutions in genes encoding surface-modifying enzymes, such as *algC* and *ykcC*. These mutations potentially alter the final protein structure while slightly modifying enzyme function. This likely enables phage resistance by hindering phage adsorption while still preserving bacterial fitness. These insights deepen our understanding of bacterial adaptation mechanisms in response to phage pressure *in vitro*. While phage therapy offers a compelling alternative or complement for pathogen control, its long-term success hinges on a comprehensive understanding of the intricate interplay between phages and bacteria, particularly considering the potential for the emergence of phage-resistant bacteria. This is critical not only for overcoming AMR bacteria in humans, animals and food but also for optimizing the biological control of plant pathogens, highlighting the diverse potential of phage therapy to shape the future of microbial management. However, relying on phages alone, even in combination with antibiotics, may not be sufficient to ensure pathogen eradication. Indeed, like antibiotic resistance, bacteria have evolved mechanisms to withstand phage attack, leading to a parallel arms race to ensure antibiotic and phage resistance. Finally, it is becoming apparent that the evolution of phage resistance may have higher fitness costs in a natural environment (*in vivo*) than in controlled laboratory conditions (*in vitro*). Because LPS have a multifaceted and pivotal role in bacterial interactions with the host, bacterial predators and competitors, and the broader microbiome, the modification of these surface decorations may involve a substantial trade-off. As a result, the emergence of phage resistance in the natural environment may be more limited.

## Acknowledgments

We thank Michael Taschner for his help with protein purification. We thank the Lausanne Genomic Technologies Facility (GTF) for the PacBio sequencing and Florian Roisne-Hamelin for his advice and help with protein visualization. This work was supported as a part of the NCCR Microbiomes, a National Centre of Competence in Research, funded by the Swiss1National Science Foundation (grant numbers 51NF40_180575 and 51NF40_225148).

## Data availability

This whole-genome shotgun project has been deposited on the EBI platform under accession number PRJEB66282. The accession number for the genome sequences of the phage-resistant variants of *Pseudomonas protegens* CHA0 are available on the EBI platform (ERS16398128; ERS16398129; ERS16398130 and ERS16398131). The raw data for bacterial growth monitoring, the output data from the variant calling analysis using Snippy and the code used are available on Zenodo (https://doi.org/10.5281/zenodo.14035382).

