## Supplementary material for "Single nucleotide switches confer bacteriophage resistance to *Pseudomonas protegen*s": Suppl. information

### Content:

*Supplementary Tables:*
**Table S1**: Bacterial strains and plasmids used in this study.
**Table S2**: Primers used in this study.
**Table S3**: SNPs detected within CDS in the natural phage-resistant variant C2.
**Table S4**: SNPs detected within CDS in the natural phage-resistant variant C4.
**Table S5**: SNPs detected within CDS in the natural phage-resistant variant C17.
**Table S6**: SNPs detected within CDS in the natural phage-resistant variant C18.

*Supplementary Figures:*
**Figure S1**: LPS profiles of the phage-resistant variants C17 and C18 and the *algC* site-directed and deletion mutants.
**Figure S2**: LPS profiles of the phage-resistant variants C17 and C18 and the *algC* site-directed and deletion mutants.

*References*

**Table S1**: Bacterial strains and plasmids used in this study.

| Strain / plasmid names | Genotype or relevant characteristics[1] | References |
|---|---|---|
| ***Pseudomonas protegens* CHA0** | | |
| CHA0 | *P. protegens* type strain; wild type; genome accession no. LS999205.1 | (Stutz 1986; Smits *et al.* 2019) |
| C2 | Natural phage ΦGP100-resistant variant of CHA0 | (Keel *et al.* 2002) |
| C4 | Natural phage ΦGP100-resistant variant of CHA0 | |
| C17 | Natural phage ΦGP100-resistant variant of CHA0 | |
| C18 | Natural phage ΦGP100-resistant variant of CHA0 | |
| Δ*algC* | CHA0 with deletion of *algC* (PPRCHA0_5962) | (Vacheron *et al.* 2023) |
| AlgC* | CHA0 producing AlgC(T428P) | This study |
| Δ*ykcC* | CHA0 with deletion of *ykcC* (PPRCHA0_1964) | This study |
| YkcC* | CHA0 producing YkcC(P229T) | This study |
| Δ*wzx* | CHA0 with the deletion of PPRCHA0_4354 | (Kupferschmied *et al.* 2016) |
| Δ*wbpL* | CHA0 with deletion of PPRCHA0_4350 | |
| Δ*obc3* | CHA0 with the deletion of the entire OBC3 gene cluster (PPRCHA0_1950 to 1958) | |
| ***Escherichia coli*** | | |
| *E. coli* S17-1/λpir | Laboratory strain | (Simon, Priefer and Pühler 1983) |
| **Plasmids** | | |
| pEMG | Expression vector; *ori*R6K, *lacZα* with two flanking I-SceI sites; Km$^R$, Ap$^R$ | (Martínez-García and de Lorenzo 2011) |
| pD(P+T)_T7N | Expression vector; T7 Promoter/terminator for *E. coli* expression, N-term. Tag; Cm$^R$ | (Taschner *et al.* 2024) |
| pD_1 | pD(P+T)_T7N containing a copy of the wild-type nucleotide sequence of *algC*; Cm$^R$ | This study |
| pD_2 | pD(P+T)_T7N containing a copy of the nucleotide sequence of the *algC variant* T428P*;* Cm$^R$ | This study |
| pME11180 | pEMG::*ykcC*; suicide plasmid containing the entire *ykcC* gene; Km$^R$ | This study |
| pME11181 | pEMG::*algC*; suicide plasmid containing the entire *algC* gene; Km$^R$ | This study |
| pME11182 | pEMG::YkcC_P229T; suicide plasmid for the replacement of P229T in YkcC; Km$^R$ | This study |
| pME11183 | pEMG::AlgC_T428P; suicide plasmid for the replacement of T428P in AlgC; Km$^R$ | This study |
| pME11184 | pEMG::Δ*ykcC*; suicide plasmid for the deletion of the *ykcC* gene; Km$^R$ | This study |
| pME11185 | pEMG::Δ*5617*; suicide plasmid for the deletion of PPRCHA0_5617; Km$^R$ | This study |
| pSW-2 | oriRK2, *xylS*, Pm::I-*sceI*; Gm$^R$ | |

[1] Ap$^R$, ampicillin resistance; Cm$^R$, chloramphenicol resistance; Gm$^R$, gentamycin resistance; Km$^R$, kanamycin resistance.

**Table S2**: Primers used in this study.

| **Name** | **Sequence 5' → 3'[1]** | **Purpose** |
|---|---|---|
| algC-F | GGAATTCTGCACCCCTTCAGATGGA | Amplification of the entire *algC* gene |
| algC-R | GGGGTACCTTCAAACAATCAAAACGGTAGTTG | Amplification of the entire *algC* gene |
| ykcC-F | GGAATTCTGCACCCCTTCAGATGGA | Amplification of the entire *ykcC* gene |
| ykcC-R | GGGGTACCTTCAAACAATCAAAACGGTAGTTG | Amplification of the entire *ykcC* gene |
| algC_T426P-F | GCCTCCAACACC**C**CCCCTGTGCTGG | Site-directed mutagenesis primers |
| algC_T426P-R | CCAGCACAGGGG**G**GGTGTTGGAGGC | Site-directed mutagenesis primers |
| ykcC_P229T-F | CCATGCGCAAAG**T**GCGTACAGAGAA | Site-directed mutagenesis primers |
| ykcC_P229T-R | TTCTCTGTACGC**A**CTTTGCGCATGG | Site-directed mutagenesis primers |
| ykcC_del_1 | GGGGTACCAGGTGTTCACCGCTTTTA | Deletion of *ykcC* |
| ykcC_del_2 | CGGGATCCTGAGCGATAATATCAACCCG | Deletion of *ykcC* |
| ykcC_del_3 | CGGGATCCCCCTGCCATTGATTCATC | Deletion of *ykcC* |
| ykcC_del_4 | ACGCGTCGACGGTATACGTCACGTGTGC | Deletion of *ykcC* |
| ykcC_check_F | CCAGAATCAGTTTTGCGTTT | Control of deletion of *ykcC* (PPRCHA0_1964) |
| ykcC_check_R | CGAGCAGGTTCTACCTGATT | Control of deletion of *ykcC* (PPRCHA0_1964) |
| algC-check-F | GCCAGCTGATCTCTTTTATC | Control of deletion of *algC* (PPRCHA0_5962) |
| algC-check-R | TTTTCGGCTTTCAATGCTTC | Control of deletion of *algC* (PPRCHA0_5962) |

[1] Site of restriction enzymes are underlined. The nucleotides involved in site-directed mutagenesis are written in bold.

**Table S3**: SNPs detected within CDS in the natural phage-resistant variant C2.

| Locus tag[1] | Position | Type of mutation[2] | CHA0 WT | C2 | Nucleotide position | Amino acid position | Product | Detected in other variants | Detected by TnSeq[3] |
|---|---|---|---|---|---|---|---|---|---|
| PPRCHA0_0474 | 543031 | del | CTGT | C | 1580/2343 | 526/780 | Putative membrane protein | - | - |
| PPRCHA0_1227 | 1399207 | del | GGTGA | G | 220/408 | 74/135 | Hypothetical protein | - | - |
| PPRCHA0_1551 | 1741067 | del | GC | G | 559/1581 | 187/526 | NADP-dependent fatty aldehyde dehydrogenase | - | - |
| PPRCHA0_1791 | 1998236 | del | GC | G | 68/2232 | 23/743 | DNA internalization competence protein ComEC/Rec2-like protein | - | - |
| PPRCHA0_1964 | 2194554 | snp | G | T | 685/978 | 229/325 | Putative glycosyltransferase YkcC | C4 | + |
| PPRCHA0_2028 | 2262454 | del | TA | T | 318/321 | 106/106 | Regulatory protein | - | - |
| PPRCHA0_2071 | 2304284 | del | CG | C | 691/861 | 231/286 | Transcriptional regulator, LysR family | C18 | - |
| PPRCHA0_2769 | 3055238 | del | TC | T | 1005/1863 | 335/620 | Outer membrane autotransporter barrel domain protein | - | - |
| PPRCHA0_2853 | 3174319 | ins | G | GT | 585/597 | 195/198 | Hypothetical protein | - | + |
| PPRCHA0_3435 | 3866379 | del | TA | T | 41/816 | 14/271 | Hypothetical protein | C4 | - |
| PPRCHA0_3662 | 4140689 | snp | C | T | 310/1176 | 104/391 | Thiolase domain protein | C4 | - |
| PPRCHA0_3692 | 4181137 | del | CCA | C | 106/276 | 36/91 | Putative secreted protein | - | - |
| PPRCHA0_3830 | 4331337 | del | CA | C | 103/195 | 35/64 | Hypothetical protein | - | - |
| PPRCHA0_4354 | 4954971 | del | CA | C | 414/1296 | 138/431 | Hypothetical protein | - | - |
| PPRCHA0_5175 | 5825570 | del | TC | T | 2175/2304 | 725/767 | PqiB family protein | - | - |
| PPRCHA0_5841 | 6542382 | del | TCTG | T | 864/966 | 288/321 | Aliphatic sulfonates ABC transporter, periplasmic sulfonate-binding protein | - | - |
| PPRCHA0_0474 | 543031 | del | CTGT | C | 1580/2343 | 526/780 | Putative membrane protein | - | - |
| PPRCHA0_1227 | 1399207 | del | GGTGA | G | 220/408 | 74/135 | Hypothetical protein | - | - |
| PPRCHA0_1551 | 1741067 | del | GC | G | 559/1581 | 187/526 | NADP-dependent fatty aldehyde dehydrogenase | - | - |
| PPRCHA0_1791 | 1998236 | del | GC | G | 68/2232 | 23/743 | DNA internalization competence protein ComEC/Rec2-like protein | - | - |

[1] Genome accession number: LS999205.1.

[2] del: deletion; ins: Insertion; sub: substitution.

[3] According to Vacheron *et al.* 2023; threshold: $Log_2$ FC > 2 and $P < 0.05$.

**Table S4**: SNPs detected within CDS in the natural phage-resistant variant C4.

| Locus tag[1] | Position | Type of mutation[2] | CHA0 WT | C4 | Nucleotide position | Amino acid position | Product | Detected in other variants | Detected by TnSeq[3] |
|---|---|---|---|---|---|---|---|---|---|
| PPRCHA0_0776 | 901430 | del | GTGGAACCT | G | 636/1446 | 212/481 | Inorganic anion transporter, SulP family | - | - |
| PPRCHA0_1209 | 1383216 | del | TG | T | 67/474 | 23/157 | 2-C-methyl-D-erythritol 2,4-cyclodiphosphate synthase | - | - |
| PPRCHA0_1216 | 1391266 | ins | G | GT | 579/2580 | 193/859 | DNA mismatch repair protein MutS | - | - |
| PPRCHA0_1280 | 1449457 | ins | G | GT | 15/1047 | 5/348 | Hypothetical protein | - | - |
| PPRCHA0_1327 | 1496986 | del | GT | G | 981/1383 | 327/460 | Transporter, major facilitator family | - | - |
| PPRCHA0_1565 | 1760560 | del | AT | A | 13/564 | 5/187 | Peptidyl-prolyl cis-trans isomerase A | - | - |
| PPRCHA0_1964 | 2194554 | snp | G | T | 685/978 | 229/325 | Putative glycosyltransferase YkcC | C2 | + |
| PPRCHA0_2099 | 2339769 | del | TG | T | 2479/2577 | 827/858 | Hypothetical protein | - | - |
| PPRCHA0_2533 | 2804712 | ins | A | AG | 849/1515 | 283/504 | Drug resistance transporter, EmrB/QacA family | - | + |
| PPRCHA0_2557 | 2826522 | del | GC | G | 68/1140 | 23/379 | Putative cytochrome P450 oxidoreductase | - | - |
| PPRCHA0_2570 | 2842067 | del | CT | C | 363/702 | 121/233 | Hypothetical protein | - | - |
| PPRCHA0_3303 | 3726100 | del | CA | C | 687/768 | 229/255 | Hydroxyacylglutathione hydrolase | - | - |
| PPRCHA0_3435 | 3866379 | del | TA | T | 41/816 | 14/271 | Hypothetical protein | C2 | - |
| PPRCHA0_3662 | 4140689 | snp | C | T | 310/1176 | 104/391 | Thiolase domain protein | C2 | - |
| PPRCHA0_4689 | 5316123 | ins | G | GT | 29/414 | 10/137 | Histidine kinase domain protein | - | - |
| PPRCHA0_5035 | 5677184 | del | CCG | C | 192/1332 | 64/443 | Sensor histidine kinase | - | - |
| PPRCHA0_0776 | 901430 | del | GTGGAACCT | G | 636/1446 | 212/481 | Inorganic anion transporter, SulP family | - | - |
| PPRCHA0_1209 | 1383216 | del | TG | T | 67/474 | 23/157 | 2-C-methyl-D-erythritol 2,4-cyclodiphosphate synthase | - | - |
| PPRCHA0_1216 | 1391266 | ins | G | GT | 579/2580 | 193/859 | DNA mismatch repair protein MutS | - | - |
| PPRCHA0_1280 | 1449457 | ins | G | GT | 15/1047 | 5/348 | Hypothetical protein | - | - |

[1] Genome accession number: LS999205.1.
[2] del: deletion; ins: insertion; sub: substitution.
[3] According to Vacheron *et al.* 2023; threshold: $Log_2$ FC > 2 and *P* < 0.05.

**Table S5**: SNPs detected within CDS in the natural phage-resistant variant C17.

| Locus tag[1] | Position | Type of mutation[2] | CHA0 WT | C17 | Nucleotide position | Amino acid position | Product | Detected in other variants | Detected by TnSeq[3] |
|---|---|---|---|---|---|---|---|---|---|
| PPRCHA0_0145 | 169392 | del | TG | T | 802/1482 | 268/493 | Efflux transporter, outer membrane factor lipoprotein, NodT family | - | - |
| PPRCHA0_0779 | 905778 | del | TC | T | 1138/1467 | 380/488 | N-acyl-D-aspartate deacylase | - | - |
| PPRCHA0_0949 | 1089977 | snp | G | C | 913/1104 | 305/367 | Oxidoreductase, FAD/FMN dependent | - | - |
| PPRCHA0_2385 | 2661991 | del | CG | C | 214/1470 | 72/489 | Hypothetical protein | C4 | - |
| PPRCHA0_2573 | 2845432 | del | TC | T | 1083/1305 | 361/434 | 2-Oxoisovalerate dehydrogenase E2 component, dihydrolipoamide acyltransferase | - | - |
| PPRCHA0_3160 | 3546111 | del | GC | G | 160/1128 | 54/375 | Atpase, AFG1 family | - | - |
| PPRCHA0_3384 | 3821536 | del | AC | A | 451/1143 | 151/380 | Glycerate kinase | - | - |
| PPRCHA0_3455 | 3880757 | del | CT | C | 9/138 | mars-45 | Hypothetical protein | C2 | - |
| PPRCHA0_3568 | 4023944 | del | AG | A | 37/810 | 13/269 | Amino acid ABC transporter, ATP-binding protein | - | - |
| PPRCHA0_3911 | 4409041 | del | GC | G | 368/753 | 123/250 | Hypothetical protein | - | - |
| PPRCHA0_4421 | 5030370 | del | CG | C | 54/786 | 18/261 | Hypothetical protein | - | - |
| PPRCHA0_4733 | 5364336 | ins | A | AC | 89/375 | 30/124 | Transcriptional regulator MvaT | - | - |
| PPRCHA0_4734 | 5365591 | ins | A | AC | 565/1428 | 189/475 | Exodeoxyribonuclease I | - | - |
| PPRCHA0_4745 | 5376849 | ins | G | GC | 633/939 | 211/312 | 2Fe-2S iron-sulfur cluster binding domain/oxidoreductase, NAD-and FAD-binding domains protein | - | - |
| PPRCHA0_4745 | 5376890 | ins | A | AT | 592/939 | 198/312 | NAD-and FAD-binding domains protein | - | - |
| PPRCHA0_5417 | 6088884 | del | GC | G | 69/1893 | 23/630 | Putative TonB-dependent outermembrane B12 receptor | - | - |
| PPRCHA0_5449 | 6123322 | del | AG | A | 863/885 | 288/294 | Hydrolase, TatD family | - | - |
| PPRCHA0_5567 | 6254238 | del | GC | G | 48/2376 | 16/791 | Quinate/shikimate dehydrogenase | - | - |
| PPRCHA0_5586 | 6277140 | del | AG | A | 21/1806 | 7/601 | Acyl-CoA dehydrogenase family protein | - | - |
| PPRCHA0_5962 | 6671173 | snp | A | C | 1282/1398 | 428/465 | Phosphomannomutase/phosphoglucomutase AlgC | C18 | + |

[1] Genome accession number: LS999205.1.
[2] del: deletion ; ins: insertion ; sub: substitution.
[3] According to Vacheron *et al.* 2023; threshold: $Log_2$ FC > 2 and *P* < 0.05.

**Table S6**: SNPs detected within CDS in the natural phage-resistant variant C18.

| Locus tag[1] | Position | Type of mutation[2] | CHA0 WT | C18 | Nucleotide position | Amino acid position | Product | Detected in other variants | Detected by TnSeq[3] |
|---|---|---|---|---|---|---|---|---|---|
| PPRCHA0_0018 | 19’321 | del | AG | A | 550/888 | 184/295 | Putative lipid A biosynthesis lauroyl acyltransferase | - | - |
| PPRCHA0_0250 | 288’360 | del | TG | T | 289/762 | 97/253 | Polar amino acid ABC transporter, ATP-binding protein | - | - |
| PPRCHA0_0480 | 549’630 | del | CT | C | 1011/1674 | 337/557 | AMP-binding domain protein | - | - |
| PPRCHA0_0862 | 992’871 | del | CT | C | 1815/2031 | 605/676 | Methyl-accepting chemotaxis protein | - | - |
| PPRCHA0_1173 | 1’341’778 | ins | G | GT | 669/2592 | 223/863 | Glycosyltransferase, group 2 family | - | - |
| PPRCHA0_2071 | 2’304’317 | del | TG | T | 727/861 | 243/286 | Transcriptional regulator, LysR family | C2 | - |
| PPRCHA0_2373 | 2’648’247 | del | CG | C | 693/1107 | 231/368 | Hypothetical protein | - | - |
| PPRCHA0_2439 | 2’711’049 | del | AC | A | 34/462 | 12/153 | Putative membrane protein | - | - |
| PPRCHA0_4628 | 5'248’148 | ins | C | CGT | 145/744 | 49/247 | NAD dependent epimerase/dehydratase family protein | - | - |
| PPRCHA0_5110 | 5'753’680 | del | CT | C | 1413/1728 | 471/575 | Hypothetical protein | - | - |
| PPRCHA0_5145 | 5'789’518 | del | GC | G | 261/1317 | 87/438 | Methyl-accepting chemotaxis transducer/sensory box protein | - | - |
| PPRCHA0_5553 | 6’234’502 | del | AC | A | 885/1275 | 295/424 | Hypothetical protein | - | - |
| PPRCHA0_5962 | 6'671’173 | sub | A | C | 1282/1398 | 428/465 | Phosphomannomutase/phosphoglucomutase AlgC | C18 | + |

[1] Genome accession number: LS999205.1.
[2] del: deletion; ins: insertion; sub: substitution.
[3] According to Vacheron *et al.* 2023; threshold: $Log_2$ FC > 2 and *P* < 0.05.

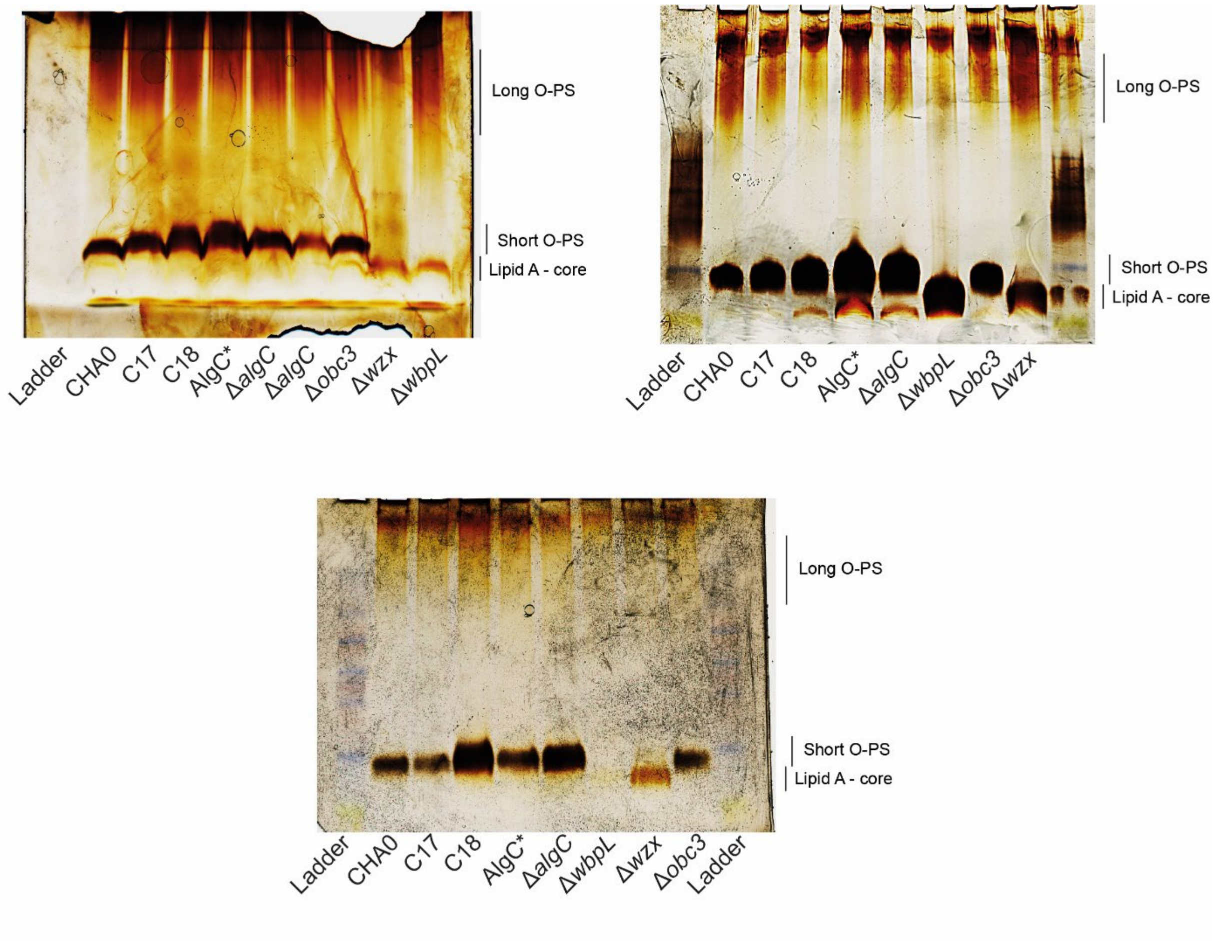


**Figure S1**: Lipopolysaccharide profiles of the phage-resistant variants C17 and C18 and the *algC* site-directed and deletion mutants.

The lipopolysaccharide (LPS) profiles of the CHA0 wild-type strain, phage-resistant variants (C17 and C18), AlgC* (site-directed mutant T229), and Δ*algC*, along with control strains Δ*wbpL*, Δ*wzx*, Δ*obc3* are displayed. SDS-PAGE was performed on LPS extracts using a 12% acrylamide gel, and the bands were visualized by silver staining. The Δ*wbpL* mutant lacks short and long O-polysaccharides (O-PS) due to a truncated core LPS, Δ*wzx* lacks short O-PS, and Δ*obc3* lacks long O-PS, as previously characterized by Kupferschmied *et al.* 2016. The bottom gel represents LPS visualized after extraction and purification using the tris-saturated phenol and diethyl ether method (Davis and Goldberg 2012).

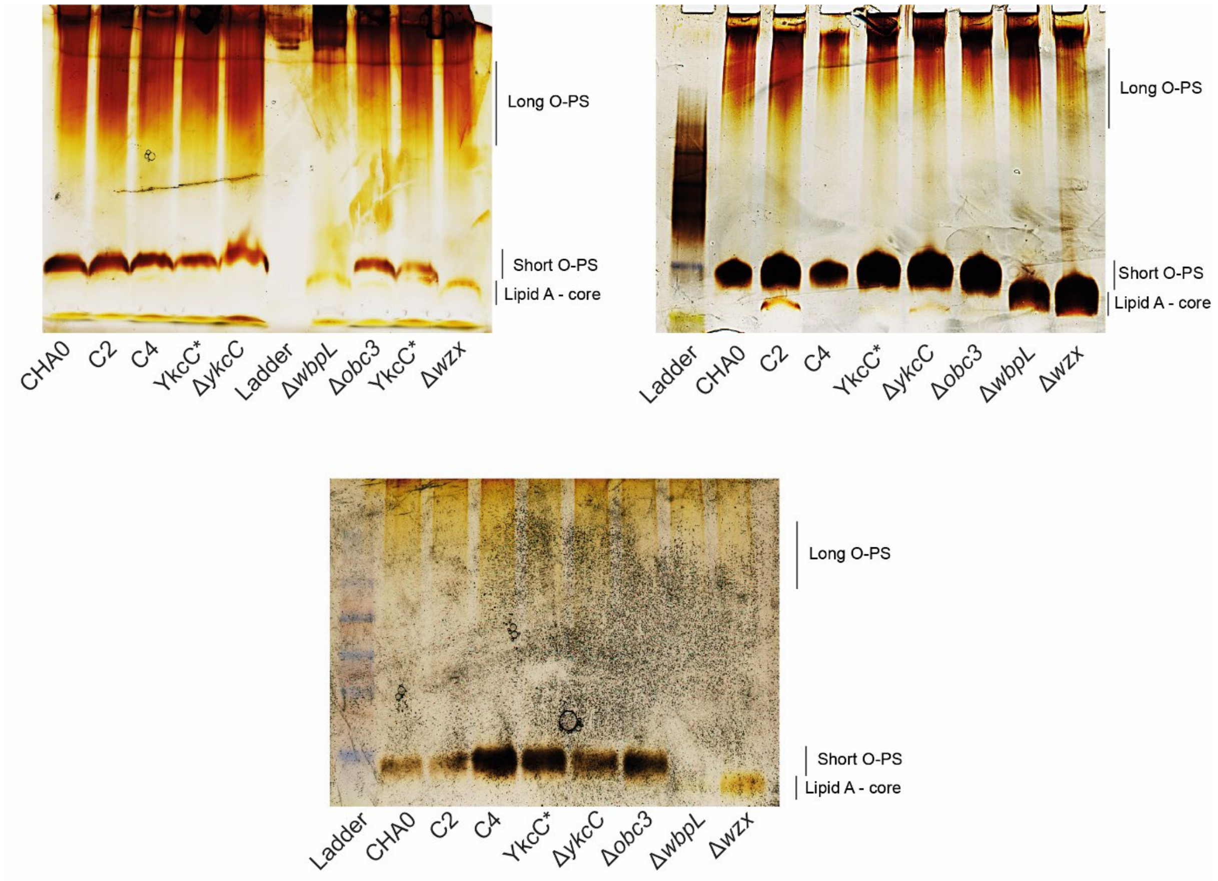


**Figure S2**: Lipopolysaccharide profiles of the phage-resistant variants C2 and C4 and the *ykcC* site-directed and deletion mutants.

The lipopolysaccharide (LPS) profiles of the CHA0 wild-type strain, phage-resistant variants (C2 and C4), YkcC* (site-directed mutant T229), and Δ*ykcC*, along with control strains Δ*wbpL*, Δ*wzx*, Δ*obc3* are displayed. SDS-PAGE was performed on LPS extracts using a 12% acrylamide gel, and the bands were visualized by silver staining. The Δ*wbpL* mutant lacks short and long O-polysaccharides (O-PS) due to a truncated core LPS, Δ*wzx* lacks short O-PS, and Δ*obc3* lacks long O-PS, as previously characterized by Kupferschmied *et al.* 2016. The bottom gel represents LPS visualized after extraction and purification using the tris-saturated phenol and diethyl ether method (Davis and Goldberg 2012).
